# Mitochondrial DNA copy number reduction via *in vitro TFAM* knockout remodels the nuclear epigenome and transcriptome

**DOI:** 10.1101/2024.01.29.577835

**Authors:** Julia Nguyen, Phyo W. Win, Tyler Shin Nagano, Elly H. Shin, Charles Newcomb, Dan E. Arking, Christina A. Castellani

**Author notes:** These authors contributed equally to this work and share first authorship (Julia Nguyen, Phyo W. Win). Corresponding author (CC).

## Abstract

Mitochondrial DNA copy number (mtDNA-CN) is associated with several age-related chronic diseases and is a predictor of all-cause mortality. Here, we examine site-specific differential nuclear DNA (nDNA) methylation and differential gene expression resulting from *in vitro* reduction of mtDNA-CN to uncover shared genes and biological pathways mediating the effect of mtDNA-CN on disease. Epigenome and transcriptome profiles were generated for three independent human embryonic kidney (HEK293T) cell lines harbouring a mitochondrial transcription factor A (*TFAM*) heterozygous knockout generated via CRISPR-Cas9, and matched control lines. We identified 4,242 differentially methylated sites, 228 differentially methylated regions, and 179 differentially expressed genes associated with mtDNA-CN. Integrated analysis uncovered 381 Gene-CpG pairs. GABA_A_ receptor genes and related pathways, the neuroactive ligand receptor interaction pathway, ABCD1/2 gene activity, and cell signalling processes were overrepresented, providing insight into the underlying biological mechanisms facilitating these associations. We also report evidence implicating chromatin state regulatory mechanisms as modulators of mtDNA-CN effect on gene expression. We demonstrate that mitochondrial DNA variation signals to the nuclear DNA epigenome and transcriptome and may lead to nuclear remodelling relevant to development, aging, and complex disease.

## Introduction

Mitochondria are cytoplasmic organelles that are essential to cell metabolism. They are central to many cellular functions such as oxidative phosphorylation, apoptotic processes, and cell differentiation via cell signalling [1]. Despite widespread evidence that mitochondrial DNA (mtDNA) mutations affect many diseases, the biological mechanisms responsible for mitochondrial dysfunction leading to age-related diseases are largely unknown. Unlike nDNA which is present in two copies per cell, mitochondria have multiple copies of mtDNA per cell (100s to 1000s). The exact number of mtDNA per cell is highly variable depending on cell type and declines with age [2]. Reductions in mtDNA-CN have been associated with reduced respiratory enzyme activity, lower expression of proteins in oxidative phosphorylation, and differences in cellular characteristics [3]. mtDNA-CN variation has been associated with many age-related diseases such as cardiovascular disease [4–7], kidney-related diseases [8,9], respiratory diseases [10,11], obesity [12], cancers [13–16], neurodegenerative diseases [17], and diabetes [18], and reduced mtDNA-CN has been associated with all-cause mortality [19]. mtDNA-CN estimates are readily obtained using DNA from blood samples, hence may serve as a potential clinically relevant biomarker of mitochondrial function and disease risk.

The nuclear epigenome consists of a collection of chemical modifications to DNA and associated proteins that may dynamically influence gene and phenotypic expressions in response to changes in the internal and external environment. DNA methylation, the most well-understood epigenetic mark, involves the addition of methyl groups on cytosine residues adjacent to guanine residues; these CG dinucleotides are called CpG sites [20]. Promoters are often located within close proximity to clusters of CpGs such that the methylation status of these CpG islands can affect gene expression by preventing interactions with transcriptional machinery [21]. Typically, transcription factors are unable to bind promoters that are heavily methylated, hence the gene cannot be transcribed [21].

Bi-directional communication between mtDNA and nDNA is essential for maintaining homeostasis, normal cell functioning, and the structural integrity of the cell [22,23]. A pioneering study reported that depletions in mtDNA levels have significant effects on nDNA methylation patterns that may contribute to tumorigenesis [24]. This initial discovery has since been supported by multiple studies that have demonstrated the link between mitochondrial DNA variation and the nuclear epigenome. Studies have identified significant associations between mtDNA-CN, CpG methylation, and gene expression in human cohorts, and *in vitro* models[25–28]. Associations between mtDNA sequence variation and the nuclear epigenome have also been observed [29–32]. Despite these associations, the mechanisms driving the relationship of mtDNA modification to the nuclear epigenome and transcriptome remain unknown.

Mitochondrial transcription factor A (*TFAM*) is a nuclear-encoded mtDNA binding protein with an essential role in mitochondrial transcription initiation and mtDNA-CN regulation, such that mtDNA-CN measures are positively correlated with *TFAM* protein level [33,34]. *TFAM* knockout has demonstrated a reduction in mtDNA-CN *in vivo* [35], with several studies reporting resulting mitochondrial dysfunction [36–39]. Studies have also revealed *TFAM* knockout to influence nDNA methylation and/or gene expression in mice [40,41]. We have used our *TFAM* knockout model to interrogate the effect of mtDNA on nDNA methylation and gene expression at specific CpGs/genes identified in an epigenome-wide association study (EWAS) of mtDNA-CN [25]. To further characterize the underlying biological mechanisms that drive the global effect of *TFAM*-induced mtDNA-CN variation on nDNA methylation, we use our *TFAM* heterozygous knockout cell line model to observe the effects of *in vitro* reduction of mtDNA-CN on the global nuclear epigenome and transcriptome profiles. Through the integration of differentially methylated and differentially expressed genes, our results provide insight into the biological mechanisms which mediate the effect of mtDNA-CN on aging-related diseases.

## Results

### *TFAM* knockout reduces mtDNA-CN and *TFAM* expression in a cell model

Stable heterozygous knockout (KO) of mitochondrial *TFAM* in HEK293T cells was generated via CRISPR-Cas9 and three resultant independent KO cell lines were confirmed by qPCR of *TFAM* DNA. Negative control cell lines (N=3) were used to benchmark *TFAM* protein expression and mtDNA-CN estimates to confirm knockout. The heterozygosity of KO lines was confirmed by qPCR of *TFAM* DNA. KO lines demonstrated a 5-fold reduction of RNA expression and an 18-fold reduction in mtDNA-CN compared to negative control lines, see Castellani et al, Figure 1 [25].

**Fig 1.**
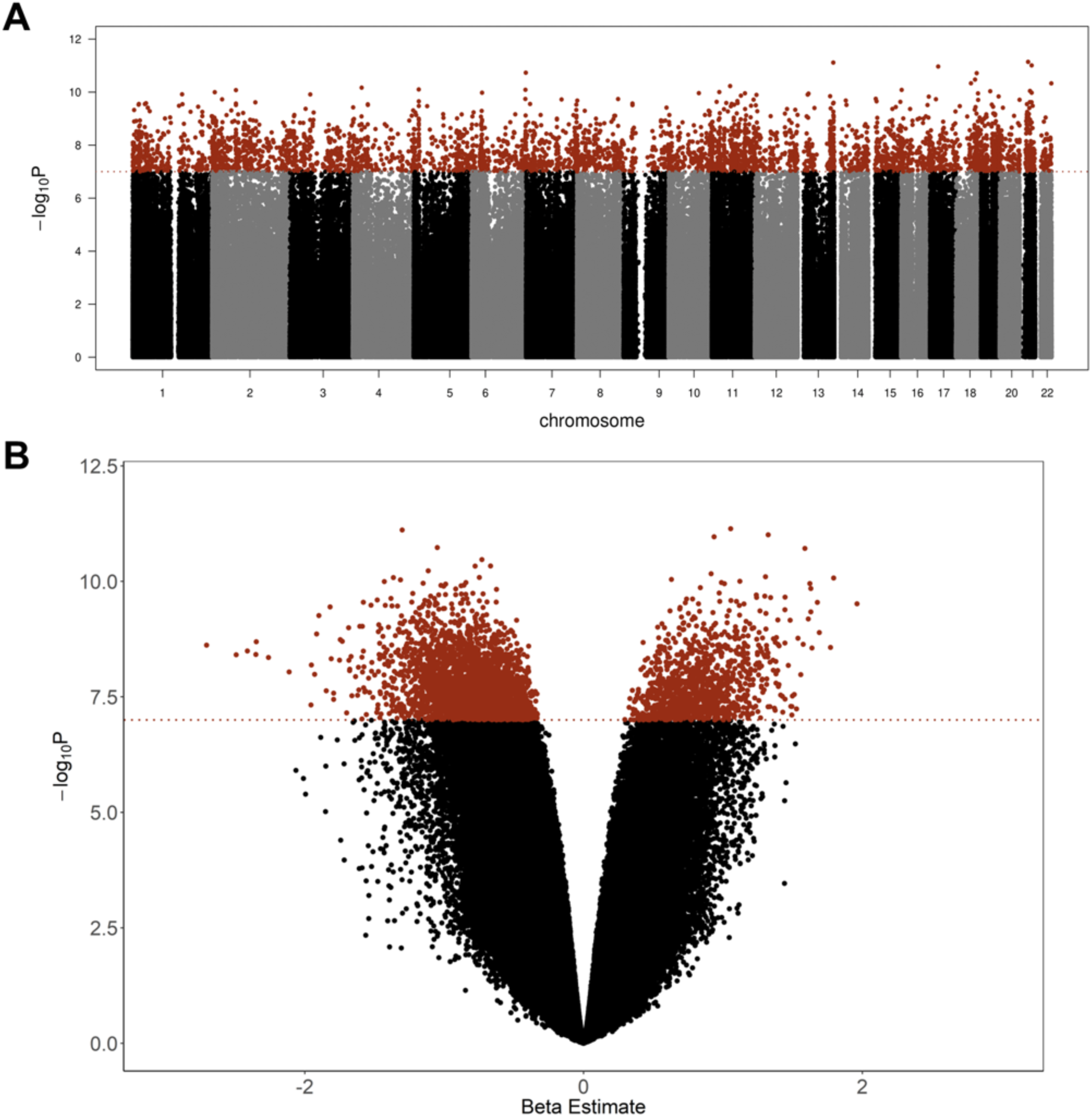
Results from differential methylation analysis. A) Manhattan plot of methylation sites across the genome. Red data points correspond to differentially methylated sites (p < 1e-07). B) Volcano plot of methylation p-values and effect sizes. Red data points correspond to differentially methylated sites (p < 1e-07).

### Site-specific nuclear methylation differences are associated with *TFAM* knockout

An EWAS using 3 *TFAM* KO cell lines and 3 normal control (NC) cell lines (all samples were run in technical duplicates) was conducted to identify site-specific differential methylation. Following normalization and quality control, 769,026 CpG sites were assessed for differential methylation between NC and KO lines. Our linear model identified 4,242 CpGs to be differentially methylated following *TFAM* KO and subsequent reduction of mtDNA-CN (p-value < 1e-7) (Fig 1A, S1 Table). No chromosomal bias was identified. However, we observed a strong bias between hyper- and hypomethylation (p-value < 2.2e-16): 1,113 / 4,242 CpGs (26%) displayed hypermethylation and 3,129 / 4,242 CpGs (74%) displayed hypomethylation in the KO groups (Fig 1B).

Previous studies using DNA extracted from blood have identified multiple CpGs that are significantly associated with mtDNA-CN [25,26]. From the study by Wang et al., 288 CpG sites were associated with mtDNA-CN (p-value < 1e-04). From that list, 2 CpG sites overlapped with the 4,242 CpG sites identified to be differentially methylated in this investigation, both with the same direction of effect (Table 1). No CpGs were found to overlap between those identified in Castellani et al. and this analysis.

**Table 1.** Differentially methylated sites that overlapped with those identified by Wang et al., 2021. Beta: beta value determined from differential methylation analysis; Wang.Beta: beta value reported in investigation from Wang et al.; P.Value: p-value from differential methylation analysis; Wang.P.Value: p-value reported in investigation from Wang et al.

| **CpG Site** | **Beta** | **Wang.Beta** | **P.Value** | **Wang.P.Value** |
| --- | --- | --- | --- | --- |
| cg01100175 | -1.1027 | -1.25092881 | 6.97E-09 | 1.45E-05 |
| cg13694927 | -0.5632 | -2.18976753 | 5.24E-08 | 8.48E-05 |

### *TFAM* knockout lines exhibit region-level nuclear methylation changes

Differential methylated region (DMR) analysis was performed to identify clusters of CpGs in the same region that were differentially methylated in *TFAM* KO lines. 228 regions were differentially methylated (Bonferroni p-value < 1.17e-5) between the *TFAM* KO and NC cells. 223 regions (97.8%) were hypomethylated in the KO groups while 5 regions (2.2%) were hypermethylated in the KO groups (S2 Table) (p-value < 2.2e-16 for hypomethylation bias). The top 4 DMRs as identified by the Fisher’s multiple comparison statistic by DMRcate exhibited reduced methylation levels in the *TFAM* KO group compared to the NC group. Biological replicates (2 per independent cell line) showed consistent clustering as can be seen by four representative plot examples (Figs 2A-D).

**Fig 2.**
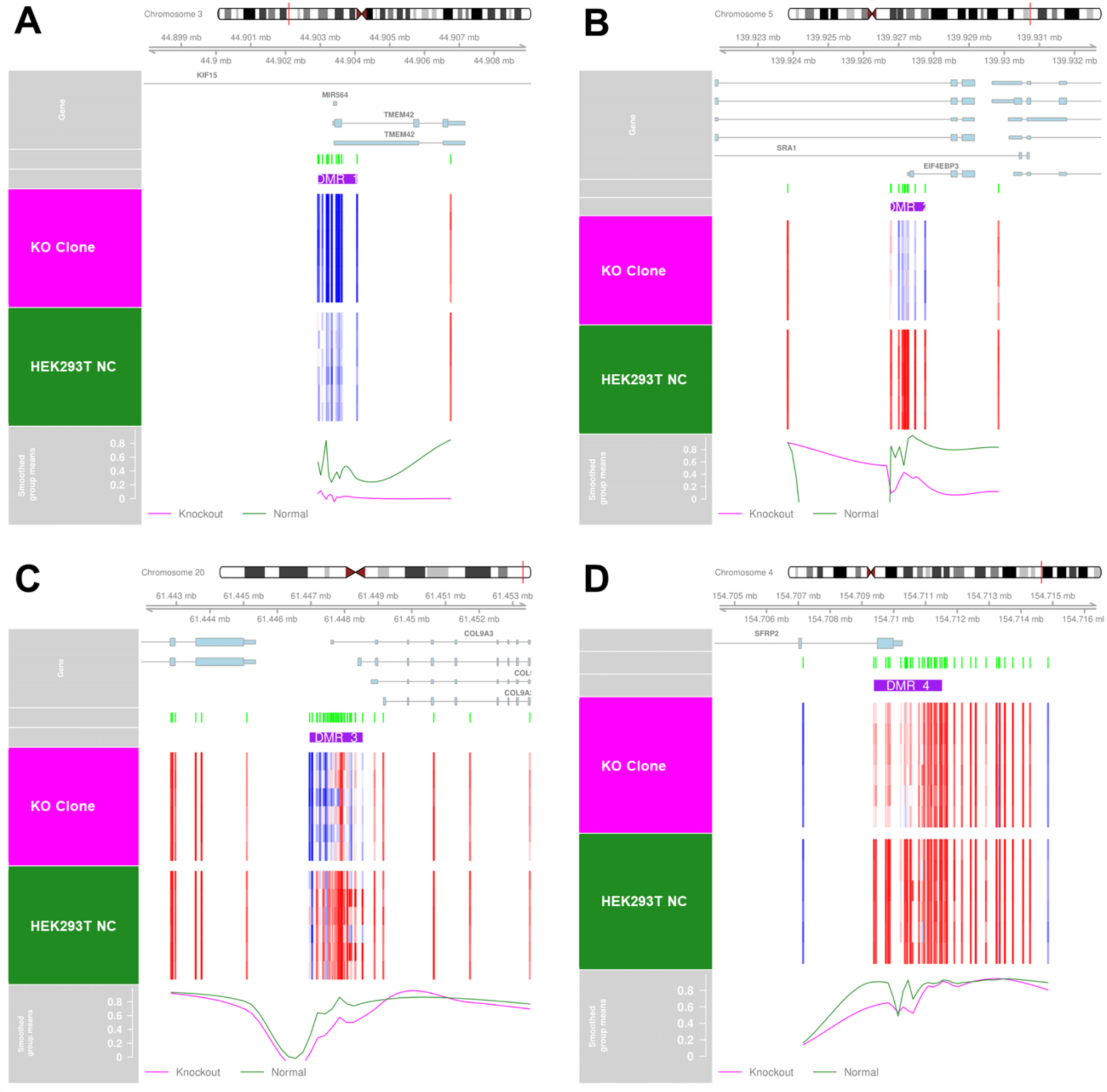
Top 4 DMRs identified. Pink represents *TFAM* KO groups, green represents NC groups. Rows represent biological replicates in a given group. A) Most significant identified DMR, located on chromosome 3. B) Second most significant identified DMR, located on chromosome 5. C) Third most significant identified DMR, located on chromosome 20. D) Fourth most significant identified DMR, located on chromosome 4. Top pink row corresponds to KO groups. Bottom forest green row corresponds to NC group. Blue panels indicate hypomethylation, red panels indicate hypermethylation.

### *TFAM* knockout leads to differential nuclear gene expression

RNA sequencing was performed to evaluate differential gene expression between our *TFAM KO* and NC cell lines, with each independent cell line analyzed in duplicate and adjusted for batch. After normalization and quality control steps were performed, 14,149 genes remained to test for differential gene expression. 179 genes were differentially expressed (Bonferroni p-value < 3.53e-6 and an absolute log fold change > 2 cut-off was determined by visualization) (S3 Table) with no chromosomal bias identified (Fig 3A). A strong bias between upregulation and downregulation of gene expression was observed (p-value < 1.8e-07): 41 / 179 genes (22.9%) were under-expressed and 138 / 179 genes (77.1%) are over-expressed in KO groups (Fig 3B).

**Fig 3.**
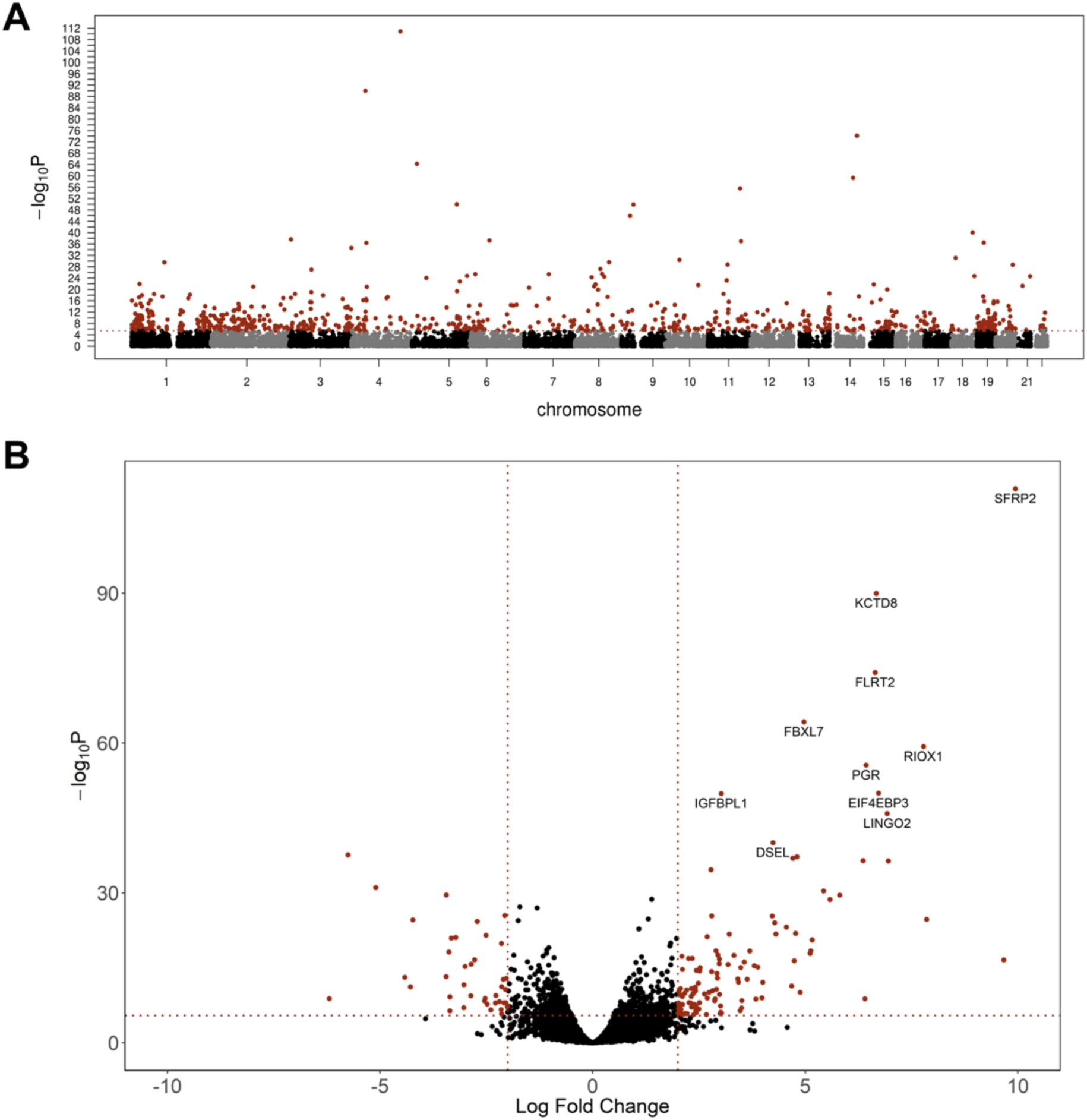
Results from differential gene expression analysis. A) Manhattan plot of gene transcripts across the genome. Red data points correspond to differentially expressed genes (p < 3.53e-6). B) Volcano plot of gene expression p-values and effect sizes. Red data points correspond to differentially methylated sites (p < 3.53e-6).

### Integration of methylation and gene expression identifies CpG-gene interactions associated with *TFAM* KO

To assess the relationship between differentially methylated sites and differentially expressed genes, we used the R package ELMER to integrate methylation and gene expression results for both direct and inverse associations between methylation at CpG sites and gene expression [42]. ELMER identified 35,347 possible Gene-CpG pairs from the 4,242 significant CpGs and the 179 DEGs. Gene-CpG pairs were filtered to include only those with differential methylation and gene expression between KO and NC (p-value < 0.001 and FDR < 0.01). Further, only CpGs within 1 Mbp from the transcriptional start site of the gene were retained. 381 gene-CpG pairs matched the filtering criteria established above, which included 65 unique genes and 353 unique CpG sites (S4 Table).

Methylation and gene expression for the top 4 unique gene-CpG pairs, prioritized by p-value and gene-CpG distance, were visualized (Fig 4). The top two identified gene-CpG pairs both belonged to genes encoding carbonic anhydrases. Both *CA2* and *CA8* genes were significantly downregulated in the gene expression analysis and further associated with hypomethylation at 2 significant CpG sites each (*CA2* with cg06015329 and cg01059952; *CA8* with cg19763925 and cg24998197) (Figs 4A-B).

**Fig 4.**
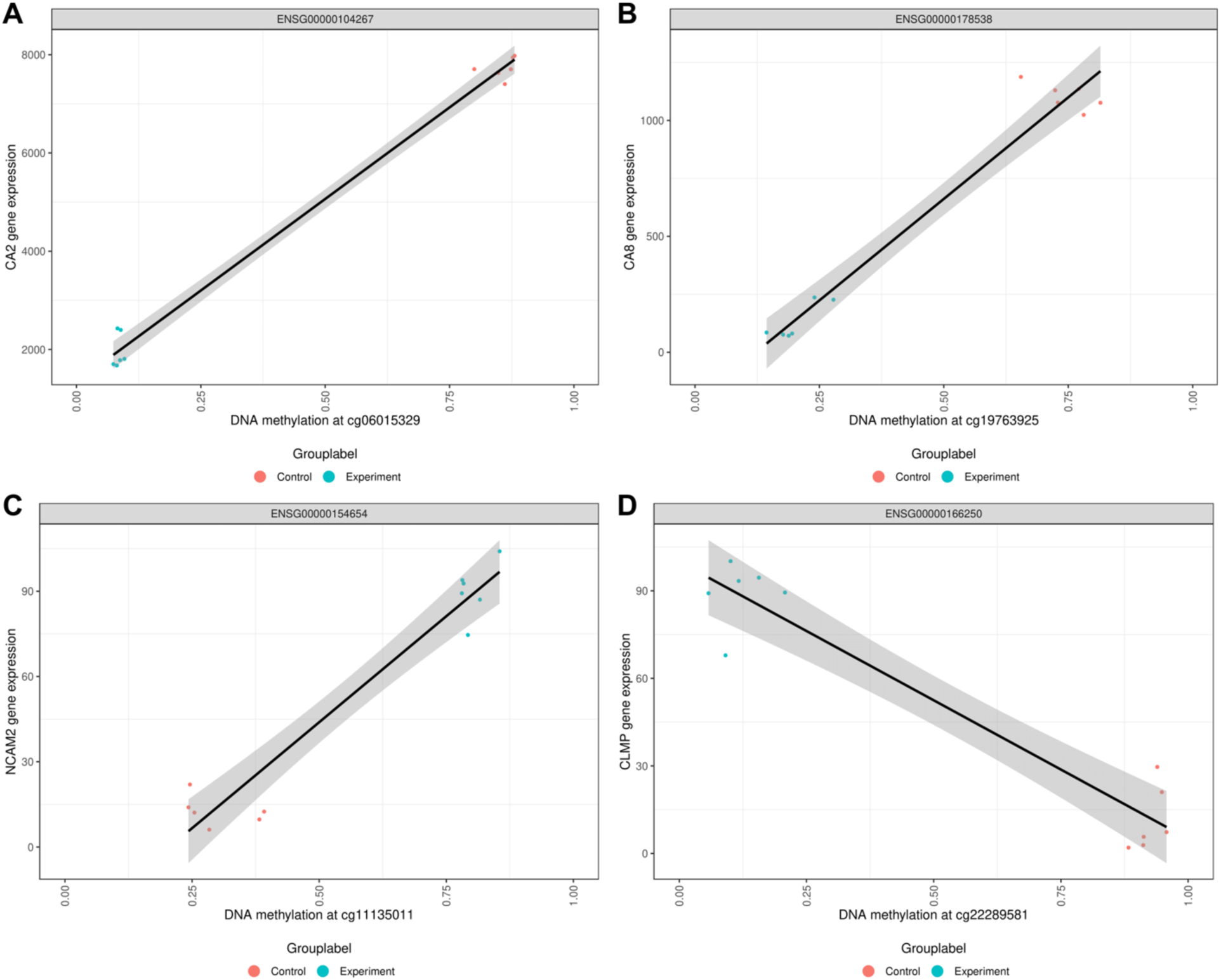
Top integrated methylation and gene expression results from ELMER analysis. A) Association between *CA2* gene expression and methylation at cg06015329, identified as the top gene-CpG pair from ELMER analysis. B) Association between *CA8* gene expression and methylation at cg19763925, identified as the second prioritized gene-CpG pair. C) Association between *NCAM2* gene expression and methylation at cg11135011, identified as the third prioritized gene-CpG pair. D) Association between *CLMP* gene expression and methylation at cg22289581, identified as the fourth prioritized gene-CpG pair. Red corresponds to NC groups, blue corresponds to *TFAM* KO groups.

### GABA and cell signaling-related genes are associated with *TFAM* knockout-induced differential methylation and gene expression

To identify biological mechanisms underlying the association between *TFAM*-induced mtDNA-CN reduction, methylation, and gene expression, we performed Gene Ontology (GO) [43,44], Kyoto Encyclopedia of Genes and Genomes (KEGG) enrichment [45–47], and Reactome analyses [48] across differentially methylated sites, DMRs, and differentially expressed genes.

The top GO terms for each of the three independent analyses include genes relating to GABA, channel activity, plasma membrane components, and signalling pathways (Figs 5A-C) (S5-7 Tables). The GO Fisher’s combined results of the methylation site with gene expression results and methylation region with gene expression results show enrichment of various pathways related to GABA, including GABA-gated chloride ion channel activity (p-value < 3.56e-10, p-value < 4.98e-10), GABA receptor activity (p-value < 3.78e-10, p-value < 4.43e-09), GABA-A receptor activity (p-value < 5.84e-10, p-value < 7.76e-09), GABA-A receptor complex (p-value < 5.84e-10, p-value < 7.76e-09), and GABA receptor complex (p-value < 2.34e-09, p-value < 1.83e-08). Genes relating to the plasma membrane were also overrepresented, including integral component of plasma membrane (p-value < 9.05e-15, p-value < 1.67e-08) and intrinsic component of plasma membrane (p-value < 9.05e-15, p-value < 2.52e-08). GO term ligand-gated anion channel activity (p-value < 2.47e-09, p-value < 6.40e-09) was also highlighted in our results.

**Fig 5.**
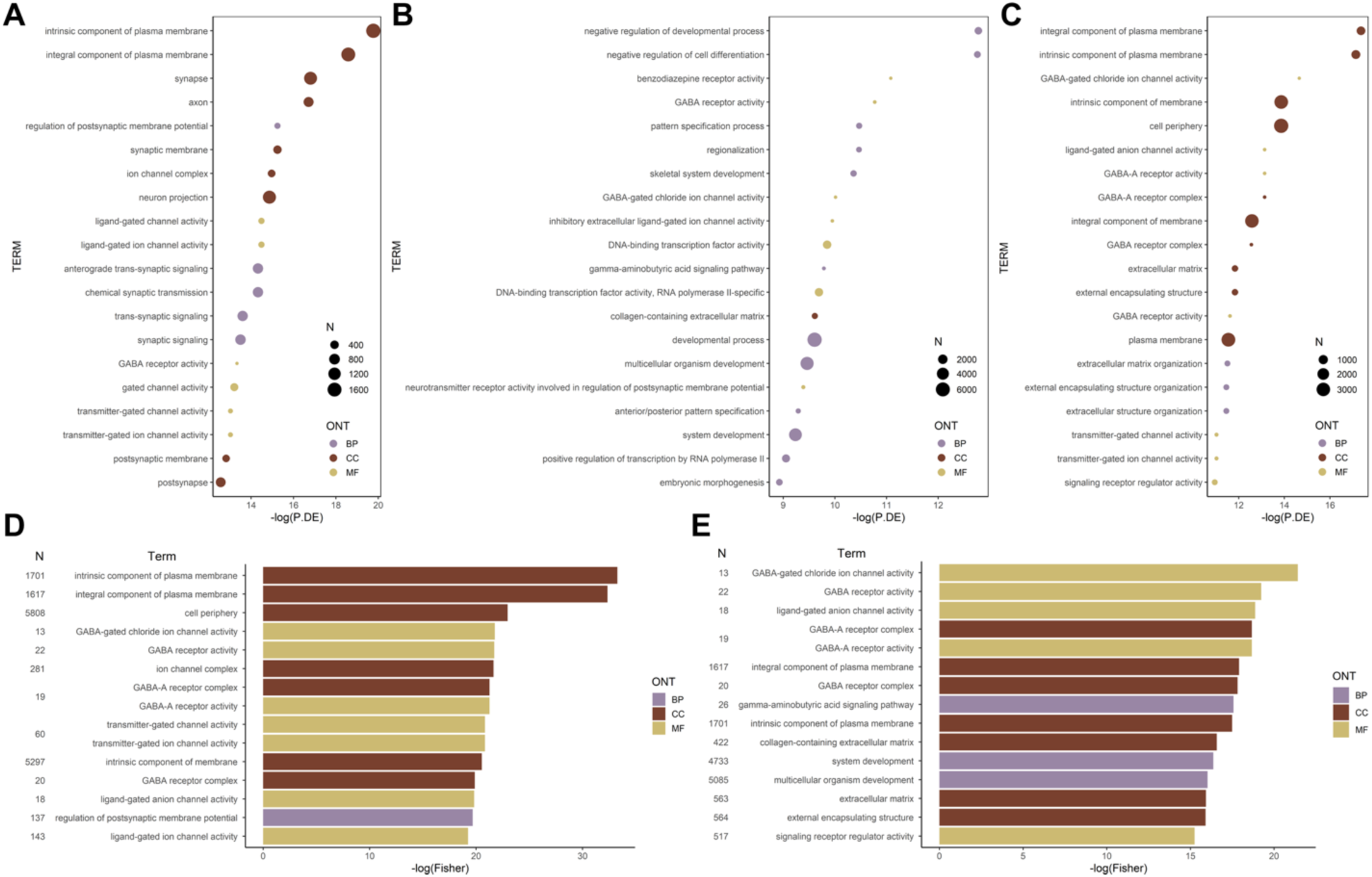
Top gene ontology (GO) terms identified. N: number of genes identified in each term; ONT: ontology; BP: biological process; CC: cellular components; MF: molecular function. A) Terms identified from differentially methylated sites (p < 1e-07). B) Terms identified from differentially methylated regions (p < 1.17e-5). C) Terms identified from differentially expressed genes (p < 3.53e-6). D) Fisher’s Combined results from methylation site and gene expression. E) Fisher’s Combined results from methylation region and gene expression.

Differentially methylated site and gene expression results were enriched for GO terms relating to channel activity, including ion channel complex (p-value < 4.00e-10), transmitter-gated ion channel activity (p-value < 9.01e-10), transmitter-gated channel activity (p-value < 9.01e-10), and ligand-gated ion channel activity (p-value < 4.40e-09) (Fig 5D, Table 2). Differentially methylated region and gene expression results were enriched for GO terms relating to signaling pathways, including gamma-aminobutyric acid signaling pathway (p-value < 2.33e-08) and signaling receptor regulator activity (p-value < 2.38e-07) (Fig 5E, Table 3). Taken together, these results implicate cell communication pathways as integral to these relationships.

**Table 2.** Overrepresented gene ontology terms identified from Fisher’s combined test of differentially methylated sites and differentially expressed genes. BP: biological process; MF: molecular feature; CC: cellular component.

| GO ID | Ontology | Term | N | DE | P.DE Meth | P.DE RNA | Fisher |
| --- | --- | --- | --- | --- | --- | --- | --- |
| GO:0031226 | CC | intrinsic component of plasma membrane | 1701 | 256 | 2.61E-09 | 3.63E-08 | 3.58E-15 |
| GO:0005887 | CC | integral component of plasma membrane | 1617 | 242 | 8.45E-09 | 2.90E-08 | 9.05E-15 |
| GO:0071944 | CC | cell periphery | 5808 | 711 | 4.10E-06 | 9.59E-07 | 1.07E-10 |
| GO:0022851 | MF | GABA-gated chloride ion channel activity | 13 | 8 | 3.17E-05 | 4.32E-07 | 3.56E-10 |
| GO:0016917 | MF | GABA receptor activity | 22 | 12 | 1.61E-06 | 9.02E-06 | 3.78E-10 |
| GO:0034702 | CC | ion channel complex | 281 | 64 | 3.17E-07 | 4.88E-05 | 4.00E-10 |
| GO:0004890 | MF | GABA-A receptor activity | 19 | 10 | 1.16E-05 | 1.97E-06 | 5.84E-10 |
| GO:1902711 | CC | GABA-A receptor complex | 19 | 10 | 1.16E-05 | 1.97E-06 | 5.84E-10 |
| GO:0022824 | MF | transmitter-gated ion channel activity | 60 | 21 | 2.21E-06 | 1.63E-05 | 9.01E-10 |
| GO:0022835 | MF | transmitter-gated channel activity | 60 | 21 | 2.21E-06 | 1.63E-05 | 9.01E-10 |
| GO:0031224 | CC | intrinsic component of membrane | 5297 | 597 | 5.09E-05 | 9.53E-07 | 1.20E-09 |
| GO:1902710 | CC | GABA receptor complex | 20 | 10 | 2.77E-05 | 3.52E-06 | 2.34E-09 |
| GO:0099095 | MF | ligand-gated anion channel activity | 18 | 9 | 5.21E-05 | 1.97E-06 | 2.47E-09 |
| GO:0060078 | BP | regulation of postsynaptic membrane potential | 137 | 40 | 2.40E-07 | 4.95E-04 | 2.84E-09 |
| GO:0015276 | MF | ligand-gated ion channel activity | 143 | 39 | 5.09E-07 | 3.70E-04 | 4.40E-09 |

BP: biological process; MF: molecular feature; CC: cellular component.

**Table 3.** Overrepresented gene ontology terms identified from Fisher’s combined test of differentially methylated regions and differentially expressed genes. BP: biological process; MF: molecular feature; CC: cellular component.

| GO ID | Ontology | Term | N | DE | P.DE_Meth | P.DE_RNA | Fisher |
| --- | --- | --- | --- | --- | --- | --- | --- |
| GO:0022851 | MF | GABA-gated chloride ion channel activity | 13 | 4 | 4.49E-05 | 4.32E-07 | 4.98E-10 |
| GO:0016917 | MF | GABA receptor activity | 22 | 5 | 2.10E-05 | 9.02E-06 | 4.43E-09 |
| GO:0099095 | MF | ligand-gated anion channel activity | 18 | 4 | 1.41E-04 | 1.97E-06 | 6.40E-09 |
| GO:0004890 | MF | GABA-A receptor activity | 19 | 4 | 1.72E-04 | 1.97E-06 | 7.76E-09 |
| GO:1902711 | CC | GABA-A receptor complex | 19 | 4 | 1.72E-04 | 1.97E-06 | 7.76E-09 |
| GO:0005887 | CC | integral component of plasma membrane | 1617 | 33 | 2.61E-02 | 2.90E-08 | 1.67E-08 |
| GO:1902710 | CC | GABA receptor complex | 20 | 4 | 2.37E-04 | 3.52E-06 | 1.83E-08 |
| GO:0007214 | BP | gamma-aminobutyric acid signaling pathway | 26 | 5 | 5.62E-05 | 1.91E-05 | 2.33E-08 |
| GO:0031226 | CC | intrinsic component of plasma membrane | 1701 | 34 | 3.22E-02 | 3.63E-08 | 2.52E-08 |
| GO:0062023 | CC | collagen-containing extracellular matrix | 422 | 18 | 6.70E-05 | 4.60E-05 | 6.34E-08 |
| GO:0048731 | BP | system development | 4733 | 101 | 9.75E-05 | 3.91E-05 | 7.77E-08 |
| GO:0007275 | BP | multicellular organism development | 5085 | 107 | 7.77E-05 | 7.10E-05 | 1.10E-07 |
| GO:0031012 | CC | extracellular matrix | 563 | 19 | 8.42E-04 | 7.24E-06 | 1.21E-07 |
| GO:0030312 | CC | external encapsulating structure | 564 | 19 | 8.61E-04 | 7.24E-06 | 1.24E-07 |
| GO:0030545 | MF | signaling receptor regulator activity | 517 | 15 | 7.07E-04 | 1.76E-05 | 2.38E-07 |

BP: biological process; MF: molecular feature; CC: cellular component.

The top KEGG pathways identified from individual enrichment analyses of differentially methylated sites, differentially methylated regions, and gene expression reveal genes involved in substance addiction, GABAergic synapse, neuroactive ligand-receptor interaction, and viral infection to be overrepresented (Figs 6A-C, S8-10 Tables). The KEGG Fisher’s combined results of both the methylation site with the gene expression results and methylation region with the gene expression results show the overrepresentation of GABAergic Synapse KEGG pathway (p-value< 2.40e-5, p-value< 2.47e-5) and signaling pathways such as the Neuroactive ligand-receptor interaction (p-value< 4.66e-8, p-value< 4.36e-7) and Retrograde Endocannabinoid signaling (p-value< 2.14e-4, p-value< 1.25e-3) (Figs 6D-E, Tables 4-5). The Cytokine-cytokine receptor interaction pathway (p-value< 3.91e-3) was also overrepresented in the combined methylation region and gene expression results.

**Fig 6.**
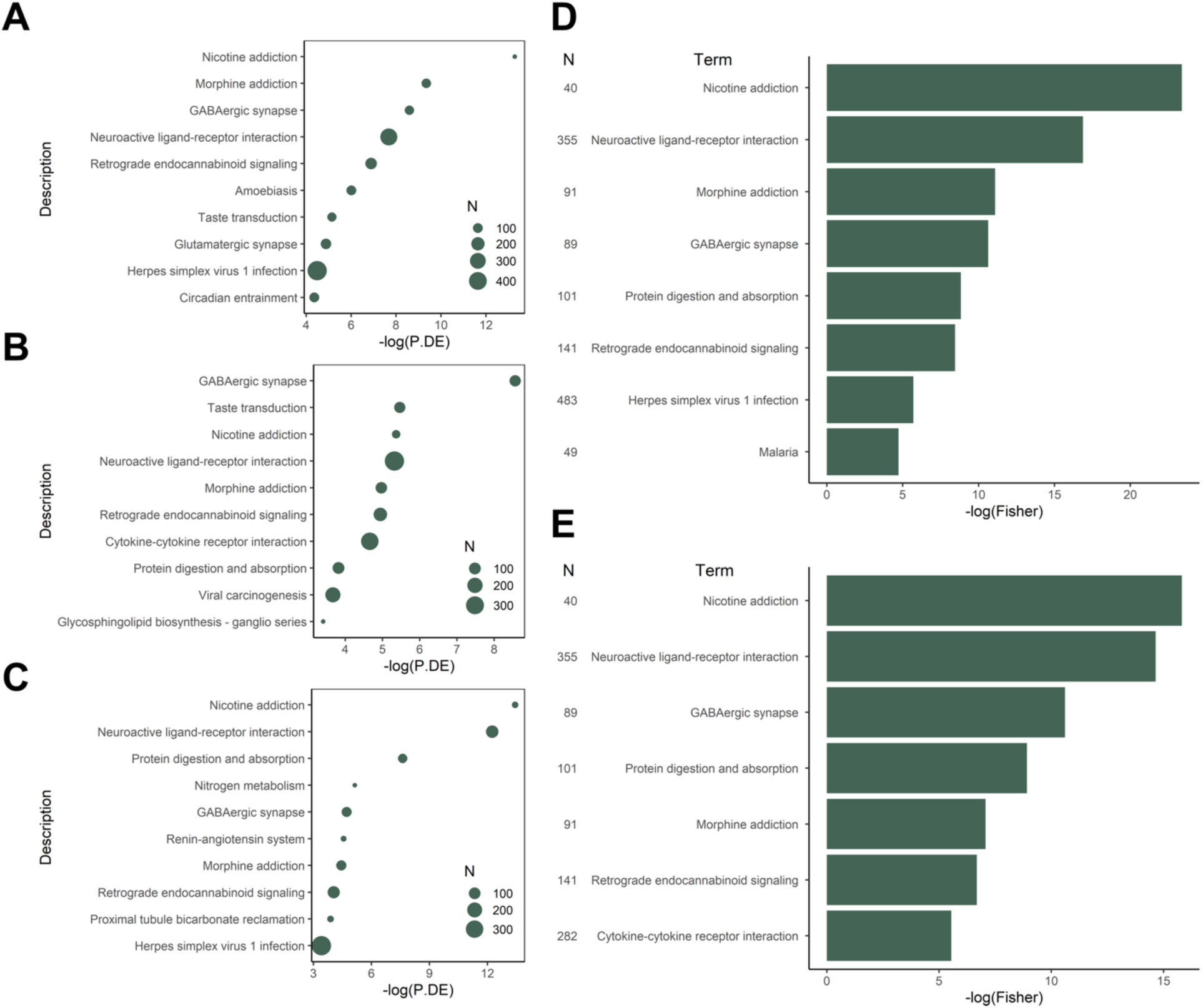
Top KEGG pathways identified. N: number of genes identified in each term. A) Pathways identified from differentially methylated sites (p < 1e-07). B) Pathways identified from differentially methylated regions (p < 1.17e-5). C) Pathways identified from differentially expressed genes (p < 3.53e-6). D) Fisher’s Combined results from methylation site and gene expression. E) Fisher’s Combined results from methylation region and gene expression.

**Table 4.** Overrepresented KEGG pathways identified from Fisher’s combined test of differentially methylated sites and differentially expressed genes.

| KEGG ID | Pathway | N | DE | P.DE_Meth | P.DE_RNA | Fisher |
| --- | --- | --- | --- | --- | --- | --- |
| path:hsa05144 | Malaria | 49 | 9 | 3.36E-02 | 3.37E-02 | 8.82E-03 |
| path:hsa05168 | Herpes simplex virus 1 infection | 483 | 55 | 1.13E-02 | 3.33E-02 | 3.34E-03 |
| path:hsa04723 | Retrograde endocannabinoid signaling | 141 | 30 | 1.02E-03 | 1.75E-02 | 2.14E-04 |
| path:hsa04974 | Protein digestion and absorption | 101 | 19 | 2.40E-02 | 4.97E-04 | 1.47E-04 |
| path:hsa04727 | GABAergic synapse | 89 | 25 | 1.85E-04 | 9.04E-03 | 2.40E-05 |
| path:hsa05032 | Morphine addiction | 91 | 27 | 8.71E-05 | 1.18E-02 | 1.52E-05 |
| path:hsa04080 | Neuroactive ligand-receptor interaction | 355 | 53 | 4.64E-04 | 4.80E-06 | 4.66E-08 |
| path:hsa05033 | Nicotine addiction | 40 | 17 | 1.70E-06 | 1.47E-06 | 6.93E-11 |

**Table 5.** Overrepresented KEGG pathways identified from Fisher’s combined test of differentially methylated regions and differentially expressed genes.

| KEGG ID | Pathway | N | DE | P.DE_Meth | P.DE_RNA | Fisher |
| --- | --- | --- | --- | --- | --- | --- |
| path:hsa04060 | Cytokine-cytokine receptor interaction | 282 | 8 | 9.45E-03 | 4.75E-02 | 3.91E-03 |
| path:hsa04723 | Retrograde endocannabinoid signaling | 141 | 7 | 7.14E-03 | 1.75E-02 | 1.25E-03 |
| path:hsa05032 | Morphine addiction | 91 | 6 | 6.94E-03 | 1.18E-02 | 8.53E-04 |
| path:hsa04974 | Protein digestion and absorption | 101 | 5 | 2.19E-02 | 4.97E-04 | 1.35E-04 |
| path:hsa04727 | GABAergic synapse | 89 | 8 | 1.91E-04 | 9.04E-03 | 2.47E-05 |
| path:hsa04080 | Neuroactive ligand-receptor interaction | 355 | 11 | 4.89E-03 | 4.80E-06 | 4.36E-07 |
| path:hsa05033 | Nicotine addiction | 40 | 4 | 4.67E-03 | 1.47E-06 | 1.36E-07 |

Top Reactome pathways identified from differentially methylated sites show results relating to channel activity, membrane potential, and the GABAergic synapse (S1 Fig, S11 Table). No overrepresented Reactome pathways were identified from gene expression results, therefore no combined analyses were performed.

Many of the identified genes converge on gene sets or pathways relating to GABA. In fact, the GABAergic synapse pathway demonstrates that the *GABA_A_* gene is hypermethylated and underexpressed in the gene expression results (S2A-C Figs). Several signaling pathways were also identified to be enriched in our datasets. This includes the Neuroactive ligand-receptor interaction pathway which was previously identified to be enriched in a human EWAS of mtDNA-CN [25]. In the neuroactive ligand-receptor interaction pathway, the *GabR* gene, a regulator of GABA, is hypermethylated in the DMR analysis and is underexpressed (S2H-I Figs), further substantiating the implication of GABA in the effects of mtDNA-CN reduction on nuclear gene expression [49].

### Differentially methylated sites are overrepresented in aging and disease-related phenotypes

The Molecular Signatures Database regulatory target gene sets were used to determine if our results were enriched for genes that targeted transcription factor binding sites [50,51]. This analysis demonstrated that 11 of 1,128 transcription factor gene sets are overrepresented in the combined methylation and gene expression results (S12 Table). Of the overrepresented gene sets, *ZNF274* is known to interact with the associated KAPI/SETDB1 histone methyltransferase complex to modulate H3K9 methylation [52].

To determine the intersection of our results with known protein-coding genes associated with mitochondrial proteins and pathways, we cross-referenced our gene set with the MitoCarta3.0 catalogue. Top results from this MitoCarta3.0 analysis reveal mitochondrial central dogma, mtRNA metabolism, small molecule transport, and calcium-related pathways to be implicated in the methylation results (S13A Table), while metabolism was the top RNA-identified pathway (S13B Table).

No overlapping pathways between methylation and gene expression results were identified in MitoCarta3.0 pathways. In addition, 5 (*ABCD1*, *ABCD2*, *PCK2*, *SARDH*, *SDSL*) of 179 significant genes from the RNA analysis are found in the MitoCarta3.0 database of mitochondrial-related genes, and all 5 genes are under-expressed (S14 Table). Though the global results are not enriched for mitochondrial genes, the *ABCD1* gene also demonstrated an association with our methylation results. This relationship showed that upregulation of *ABCD1* was associated with both hypermethylation and hypomethylation of nine significant CpG sites (cg00207916, cg01022618, cg01083397, cg05365121, cg15373098, cg19514407, cg20844535, cg24324483, and cg26149887).

### *TFAM* knockout differentially methylated sites are overrepresented for cancer and autoimmune disease-related CpGs

To explore if *TFAM* knockout-induced differentially methylated CpGs sites were associated with disease-related sites, we leveraged the MRC-IEU EWAS catalog. This EWAS catalog is a curated database of CpG sites that have been found to be associated with traits/exposures across over 2,000 EWAS publications [53] From our list of differentially methylated sites, we prioritized independent CpGs within a range of 1Mbp by selecting for the most significant CpG in each window. For both global (all-disease) and individual disease terms, a permutation test was performed on the EWAS catalog terms to identify if there was enrichment for our differentially methylated sites for each disease term. Of the 4,242 differentially methylated sites identified in this study, we identified 785 independent CpGs that overlap with those in the EWAS catalog (S15 Table). Although these CpGs were not enriched for overall disease terms (p-value = 0.347), they were enriched for cancer-related CpGs (115 CpGs, permutedp-value = 0.001) and CpGs related to autoimmune diseases (93 CpGs, permuted p-value = 0.011) (S3 Fig).

### Epigenetic regulation of non-coding regions may be modulating mtDNA-nuclear crosstalk

Genomic region enrichment testing was performed to determine if identified CpG, gene, and chromatin state regions are overrepresented in our datasets. We discovered significant overrepresentation (FDR < 0.01) of intergenic regions, enhancers, zinc finger proteins (ZNF) genes and repeats, heterochromatin, bivalent TSS, flanking bivalent TSS/Enhancer, bivalent enhancer, repressed polyComb, weak repressed polyComb and quiescent/low group in the methylation dataset (Fig 7, S16 Table). Conversely, the 5’UTR was under-enriched (FDR < 0.01) and overrepresentation of intergenic regions, different enhancer types, heterochromatin and repressed PolyComb regions was observed, implicating epigenetic marks in non-coding regions of the heterochromatic and enhancer region to be overrepresented in the overall methylation analysis.

**Fig 7.**
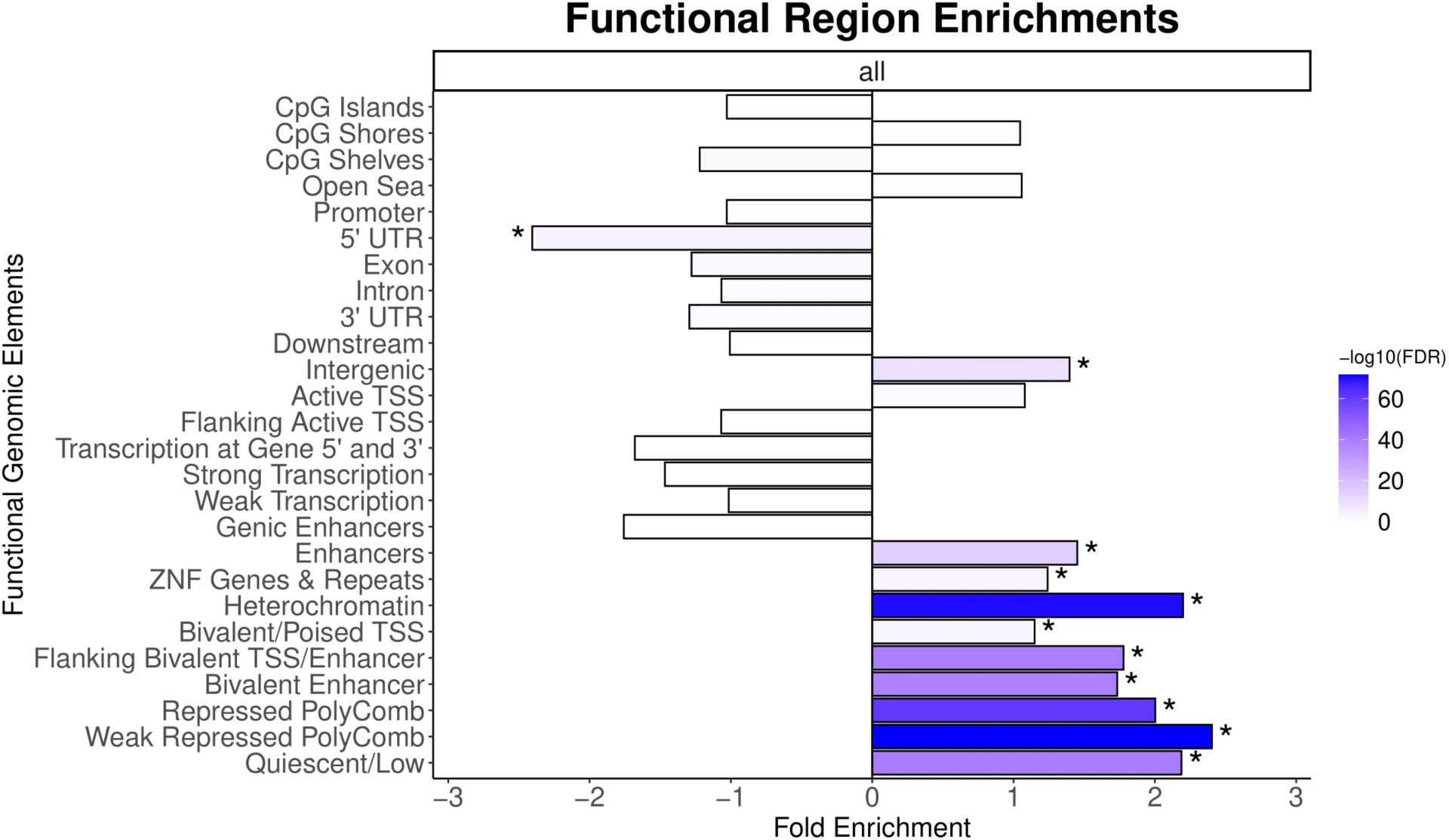
Enrichment of CpG, gene, and chromatin state regions in methylation data. FDR: false discovery rate; TSS: transcription start site; ZNF: zinc-finger. *indicates significant enrichment (FDR < 0.01).

## Discussion

To determine the genes, regulatory sites, regions, and biological pathways associated with mtDNA-CN variation, we analyzed DNA methylation data from methylation microarrays and RNA sequencing data generated from cell lines with a heterozygous knockout of the *TFAM* gene. We report widespread differential methylation and differential gene expression in *TFAM* KO samples with reduced mtDNA-CN. The results specifically implicate GABA receptors, neuroactive ligand-receptor interaction pathways, the *ABCD1* gene, cellular signaling, regulation of carbonic anhydrases, and non-coding regulation as candidates for methylation-mediated mitochondrial gene regulation.

### GABA receptor and neuroactive ligand-related processes and pathways are associated with mtDNA-CN-induced nuclear genome modifications

The combined methylation and gene expression enrichment analyses identified numerous GABA-associated pathways, including GABA-gated chloride ion channel activity, GABA receptors, and GABAergic synapse. Additionally, Nicotine Addiction, Neuroactive Ligand Receptor Interaction, Morphine Addiction, GABAergic Synapse, Protein Digestion and Absorption and Retrograde Endocannabinoid Signaling pathways are overrepresented, and these pathways are all related to cell-signaling and the nervous system.

A previous study had found an association between the neuroactive ligand receptor interaction pathway and mtDNA-CN following a multi-ethnic EWAS meta-analysis across multiple independent human cohorts [25]. Together, these studies provide further evidence that these changes are likely mediated through mtDNA-CN and that similar mechanisms may be involved. In addition, the cytokine-cytokine receptor interaction pathway was also overrepresented in the methylation site-specific and gene expression results. This finding aligns with the discovery of the role of mtDNA variability in modulating cytokine and cytokine receptor expression as part of cellular stress responses [54,55]. Both KEGG-identified pathways are classified under the Environmental Information Processing and Signaling Molecules and Interaction KEGG category, indicating the importance of cellular signaling in transmitting the effect of mtDNA-CN variability on nDNA methylation and gene expression.

The GABAergic synapse pathway was overrepresented in the integrated KEGG results, where gene expression results were combined with both differentially methylated sites and regions. Additionally, GO results revealed enrichment for GABA and GABA_A_ related gene groups in our dataset. The GABA_A_ receptor is comprised of four subunit genes: *GABRB1*, *GABRB3*, *GABRE*, and *GABRG1* [56]. In our study, all four GABA receptor genes were significantly downregulated in the *TFAM* KO samples relative to the NC samples. In addition, integration analysis found the *GABRB1* gene to be associated with site-specific methylation at 13 CpGs. The GABA_A_ receptor functions as a ligand-gated ion channel that allows for fast inhibitory synaptic transmission [56]. Reduced expression of the GABA_A_ receptor genes reduces available receptors for the binding of the GABA neurotransmitter, leading to cells with over-excitation of their terminal neurons. Additionally, the GABA neurotransmitter plays an important role in a variety of cellular processes such as suppressing inflammatory immune responses [57] and binding to islet alphas-cells to inhibit the secretion of glucagon [58]. Further, a mouse model study demonstrated mice lacking GABA-synthesizing genes die at birth despite no visible structural differences in their brains, suggesting further associated functions of GABA in neurodevelopment and animal physiology [56]. Taken together, reduced mtDNA-CN in *TFAM* KO samples appears to be associated with GABA_A_ receptor genes, which may lead to abnormal GABAergic signaling. This may provide a mechanism in which mtDNA-CN is related to age-related chronic diseases.

### Mitochondrial-related genes are under-expressed in *TFAM KO* and associated with mtDNA-CN-induced DNA methylation and gene expression with implications for disease

The family of carbonic anhydrase (CA) enzymes are found in the mitochondria across tissue types where they elicit a variety of physiological functions [59]. CAII, encoded by the *CA2* gene, is involved in several pathological conditions including neurodegeneration, glaucoma, epilepsy, and altitude sickness, which has led to growing interest in the clinical use of carbonic anhydrase inhibitors [60–62]. Patients with osteopetrosis and renal tubular acidosis were discovered to carry mutations resulting in the downregulation of CAII [63]. These manifestations are also observed in inherited autosomal recessive CAII deficiency syndrome [64]. CAVIII, encoded by the *CA8* gene, is another carbonic anhydrase isoform that is downregulated in renal cell carcinoma [65] and congenital ataxia [66]. Our results found both *CA2* and *CA8* to be downregulated following *TFAM* KO and associated with hypomethylation at 2 unique CpG sites each (4 CpG sites total) (S4 Table).

The *ABCD1* gene functions as an ATP-binding cassette transporter which helps move very long chain fatty acid-CoA from the cytosol to the peroxisome [67,68]. Additionally, the *ABCD1* gene also regulates mitochondrial-related functions such as oxidative phosphorylation and fatty acid synthesis [67,68]. The X-linked adrenoleukodystrophy genetic peroxisome disease has been linked to mutations in the *ABCD1* gene, which affects the nervous system and kidney [69,70]. Our gene expression results identify the *ABCD1* gene and it is also included in the MitoCarta3.0 database. The *ABCD1* gene is also associated with 9 significant CpG sites from our dataset (S4 Table). Furthermore, *ABCD1,* and its paralog *ABCD2*, are both underexpressed in our analysis and only the metabolism pathway overlapped between the MitoCarta3.0 database and the overrepresented pathways from the gene expression analysis. mtDNA-CN reduction following *TFAM* KO may lead to *ABCD1* hypermethylation, resulting in decreased expression and consequently altered energy metabolism. Further investigations on a potential association between mtDNA-CN alterations and *ABCD1*-related diseases such as X-linked adrenoleukodystrophy is warranted [71].

### mtDNA-CN reduction is related to the enrichment of non-coding chromatin regions

Enrichment in chromatin state regions such as enhancers and repressed Polycomb was identified in our analyses. These findings may suggest the existence of interaction between mtDNA-CN and the machinery regulating chromatin states as another avenue of mitochondrial-nuclear crosstalk, with potential implications on disease etiology. Further investigations of the associations between mtDNA-CN and chromatin states by leveraging ATAC-seq or ChIP-seq is warranted.

No CpG-related regions (islands, shores, shelves, or open sea) were overrepresented, which is consistent with a similar previous EWAS study on mtDNA-CN [26]. The enrichment for intergenic regions in our dataset follows another mtDNA-CN-modifying investigation that found DMRs overrepresented in intragenic regions, with the majority found in introns [72]. These findings further suggest gene expression control is being regulated through non-coding regions.

### mtDNA-CN remodels the nuclear epigenome and transcriptome

Our results, alongside emerging evidence on the association between mtDNA variability, the nuclear epigenome, and nuclear gene expression, reinforce the complex interplay between the mitochondria and the nuclear genome [73–75]. Our results demonstrate a reduction in mtDNA-CN and widespread changes to nDNA methylation and gene expression following *TFAM* knockout. These dynamics have also been found to be influenced by environmental stimuli [76]. It has been proposed that these environmental stimuli affect the bidirectional cross-talk between the mitochondrial and nuclear genome through several key mitochondrial metabolites [77]. Here, we identify biological pathways in relation to these associations and highlight key targets for further research. With advancements in both mitochondrial genome editing and mitochondrial transplantation as potential avenues to revolutionize the prevention and treatment of mitochondrial-related diseases, these findings provide insight on anticipated effects to the nuclear epigenome and nuclear genome expression as a result of mtDNA modifications.

Some limitations of these results include that *TFAM* KO cells have resulted in a more substantial reduction in mtDNA-CN compared to what is generally observed in human patients and should therefore be interpreted with caution. Additionally, the *TFAM* gene may also independently impact other pathways in addition to regulating mtDNA-CN and therefore could impact nDNA methylation and gene expression through unknown biological mechanisms. Furthermore, the kidney-derived HEK293T cell lines in which the *TFAM* KO and NC cell lines were established may have specific baseline methylation patterns not found in other cell types. Therefore, the findings of this study are not expected to be generalizable to all cell types.

## Conclusion

In conclusion, we report the impact of reduction of *TFAM* on site-specific differential DNA methylation, differentially methylated regions, and differential gene expression. Overrepresentation of GABA_A_ receptor genes, the neuroactive ligand receptor interaction pathway, the *ABCD1/2* genes, regulation of carbonic anhydrases, and cell signalling processes were identified. Furthermore, enrichment of functional genomic regions demonstrated that chromatin states such as enhancers and heterochromatin are overrepresented in the methylation results, implying that mtDNA-CN may also be impacting gene expression via chromatin states. The results indicate that mitochondrial DNA variation influences the nuclear DNA epigenome and transcriptome and has the potential to contribute to processes related to development, aging, and complex disease risk.

## Materials and Methods

### *TFAM* Knockout Cell Line Model

The experimental models generated in this study were modified HEK293T cell lines with a heterozygous knockout of the *TFAM* generated via CRISPR-Cas9 using the Origene *TFAM* – Human Gene Knockout Kit following the manufacturer’s protocol. The sgRNA guide sequence used was GCGTTTCTCCGAAGCATGTG. Turbofectin 8.0 was used to perform lipofection, and puromycin was used for selection. The knockout procedure was performed independently for each of the 3 KO cell lines. 3 NC cell lines were generated using the same original cell line performing all identical steps but without performing the knockout. This resulted in 6 independent samples (3 KO and 3 NC). For all downstream analyses, each independent sample was run in a minimum of technical duplicates. qPCR was used to measure mtDNA-CN as an estimate relative to nuclear DNA copy number and to confirm the number of copies of *TFAM* in cell lines as well as to confirm the number of copies of *TFAM* DNA. A Western Blot was performed to demonstrate the reduction in *TFAM* protein in the KO lines with Tubulin as the control. *TFAM* gene expression was also measured using qPCR. Gene expression was determined using the double delta cycle threshold method.

### DNA/RNA Sequencing Data Generation

DNA was extracted using the AllPrep DNA/RNA Mini Kit following manufacturer’s protocol and quantified using a Nanodrop 1000. Following bisulfite conversion, DNA strands were hybridized to the Illumina Infinium EPIC BeadChip to determine the DNA methylation profile for over 850,000 CpGs in the human genome. All samples (3 KO and 3 NC) were run on the EPIC BeadChip at two separate time points which resulted in 12 individual runs represented by 12 raw .idat files (6 KO and 6 NC).

RNA extraction, quantification, quality control, library preparation, and sequencing were performed using the Qiagen AllPrep DNA/RNA Mini Kit, Qubit 2.0 Fluorometer, and Illumina’s TruSeq Stranded Total RNA kit. Total RNA was converted to cDNA, and end repair was performed. Fragments were ligated and size selected. The Illumina HiSeq 2500 instrument was used to perform paired-end 150 bp sequencing to generate gene expression profiles. Each of the 6 samples (3 KO and 3 NC) were sequenced twice which resulted in 12 FASTQ files.

### nDNA Methylation Analysis of Cell Lines

The minfi package was used for EPIC BeadChip analysis and quality control to remove poor quality probes [78]. The data was normalized using functional normalization [79] and poor quality probes were removed if the probe had a detection p-value > 0.01. Cross-reactive probes were identified and omitted [80], as well as probes with known SNPs at the CpG site. Additionally, CpGs with greater than a 0.15 difference in mean beta value between the first and second runs for the NC cell lines were removed. This step accounted for measurement variation inherent in the EPIC BeadChip.

Differentially methylated sites were determined using a linear model with mtDNA-CN as the independent variable and methylation values as the dependent variable using both DMPFinder in minfi and a linear mixed model, with sample ID included as a random variable, where possible[78]. A significance threshold of p-value < 1e-7 was used to determine differentially methylated sites as previously demonstrated to be appropriate for EWAS analyses [81–84].

DMRs were determined via linear modelling using the R package DMRcate with treatment status (KO vs NC) as the independent variable and methylation beta value as the dependent variable [85]. The batch (1^st^ and 2^nd^ run) was included as a fixed effect. A significance threshold of p-value < 1e-7 was used to determine differentially methylated sites prior to DMR identification. Regions were defined as having a minimum of 10 CpGs with at most 1000 bps between CpGs within the region. Additionally, regions were removed if the region had a mean beta value difference < 0.05 between KO and NC. A significance threshold of p-value < 1.17e-5 was used to identify significant DMRs which reflects Bonferroni correction for multiple testing (4,259 tests).

### RNA-Sequencing Analysis of Cell Lines

Kallisto was used to pseudo-align the compressed FASTQ files to the Genome Reference Consortium Human Build 37 to generate transcript level counts [86]. To account for uncertainties in transcript counts, a total of 100 bootstraps were performed in Kallisto. Transcript level counts were converted to gene level counts using the transcript count of the isoform with the highest expression.

Normalization, quality control, and differential expression analysis were performed using the EdgeR package [87]. The data was normalized to scale for effective library size of samples using the trimmed mean of M-values (TMM) [88]. Genes with less than 15 reads across all samples were removed due to low expression. In addition, genes with a sum of counts per million (CPM) across samples < 6 were also discarded from the analysis. Samples for a gene were included in the sum of CPM if the CPM for the sample was > 0.6.

Gene expression was modelled using a weighted likelihood empirical Bayes method as described by Chen et al. [89]. Differentially expressed genes were determined using a likelihood ratio test between the null and the observed model, with the batch (1^st^ and 2^nd^ run) adjusted for as a fixed effect. Genes with an absolute log fold-change (logFC) < 2 were removed. A significance threshold of p-value < 3.53e-6 was used to determine differentially expressed genes which reflects Bonferroni correction for multiple testing (14,149 tests).

### Integration of Methylation and Gene Expression

The ELMERv2 [42] package was used to integrate differential methylation and differential gene expression to find associations between methylation and gene expression of nearby genes. Associations with any direction of effect were considered. The supervised approach was used such that methylated and unmethylated groups were split into KO and NC labels. Significant CpGs from differentially methylated site analysis and significant genes from differential expression analysis were used as the methylation and the gene expression data inputs respectively.

Each CpG site was mapped to the nearest 20 genes (10 upstream and 10 downstream) in the genome. Each Gene-CpG pair was tested using a one-sided Mann-Whitney U test to check for methylation differences between KO and NC (p-value < 0.001). Genes in Gene-CpG pairs were tested for differential gene expression between KO and NC using the Student’s t-test (p-value < 0.001). Following the same parameters to previous studies, Gene-CpG pairs were removed from the analysis if the CpG was greater than 1 Mbp from the transcriptional start site of the gene using the ENSEMBL [90] gene level annotations [25,91]. A significance threshold of false discovery rate (FDR) < 0.01 was used to determine significant Gene-CpG pairs.

### Enrichment Analyses

Enrichment analyses were performed on the differential methylation sites/regions and differential gene expression results with all tested sites, regions, or genes as background, respectively. We leveraged the following databases to uncover physiological mechanisms overrepresented in our results: The Gene Ontology (GO) [43,44] database for functional gene groups, the Kyoto Encyclopedia of Genes and Genomes (KEGG) [45–47] for biological pathways, the Reactome Pathway and transcription factor gene sets from the Molecular Signatures Database (MSigDB) [50,51], and the MitoCarta3.0 [92] database for mitochondrial related genes and pathways.

GO and KEGG over-enrichment analysis was performed on the significant methylation sites and methylation region results using missMethyl [93]. missMethyl is particularly well suited for this as it accounts for the bias introduced from unbalanced CpG-gene mapping. GO and KEGG functional enrichment analyses was performed on significant genes from RNA-sequencing using limma [94]. The top KEGG pathways were visualized using Pathview to map up/down-regulation of genes within pathways [95]. Reactome pathway functional enrichment [48] was performed on differentially methylated sites using the MethylGSA package [96] using p-value < 1e-07 and differentially expressed genes using the ReactomePA package [97].

MSigDB transcription factor binding sites regulatory target gene sets and MitoCarta3.0 mammalian mitochondrial protein sets and pathways were used to perform over-enrichment analysis on significant methylation and gene expression results. Overrepresentation of transcription factor genes was determined by using the Student’s t-test on significant genes in the gene set against the genes that were not present in the gene set. A significance threshold of p-value < 0.05 was used. In addition, the significant genes from the RNA analysis were tested for overlaps with genes in MitoCarta3.0 using a Chi-squared test (p<0.05).

For the GO, KEGG, and transcription factor enrichments, the methylation and gene expression results were combined using the Fisher’s combined probability test to combine p-values [98]. Fisher’s combined probability test could not be applied to Reactome results due to the lack of gene expression results following p-value filtering. The methylation site and the methylation region results were combined separately with the gene expression results using this method. Terms/groups with p-value > 0.05 in either the methylation or gene expression results were removed before combining p-values.

The EWAS Catalog was used to identify any differentially methylated CpG sites that have been previously reported in the literature to be associated with known phenotypes. CpG sites and their associated phenotypic traits were obtained from the EWAS Catalog website in June 2023. Traits were further collapsed into broader disease categories. Pruning was first performed to select for the most significant CpG per 1Mbp from our significantly differentially methylated probes and separately for all probes. To assess for enrichment in overlapping CpGs with given specific disease terms, 1,243 CpGs were randomly selected from the EWAS Catalog (representing the total number of pruned CpGs from our *TFAM* knockout differently methylated probes). Further, the overlap between the randomized set and each disease category was recorded. This was repeated for 1,000 permutations with the p-value measured as the percentage of the number of permutations in which the number of overlapping CpGs was greater than the observed number of overlapping CpGs.

Enrichment of genomic regions was performed using the DMRichR [99] package to test for overrepresented CpG categories (CpG islands, shores, shelves, and open sea) and gene regions (promoter, UTR, exon, intron, downstream, intergenic, active TSS, transcription, enhancers, ZNF genes and repeats, heterochromatin, PolyComb, quiescent/low). The enrichment testing for overrepresented CpG and gene regions used a Fisher’s Exact test at a significance threshold of FDR < 0.01. The DMRichR package also includes a wrapper for LOLA [100] which tests for enrichment of 15 chromatin states from ChromHMM [101]. Enrichment testing for chromatin states used a Fisher’s Exact test, and a significance threshold of FDR < 0.01 was used.

## Acknowledgements

C.A.C., P.W.W. and E.H.S. are supported by the Department of Pathology and Laboratory Medicine at Western University and The Children’s Health Research Institute (CHRI). P.W.W. is supported by the Ontario Graduate Scholarship (OGS). D.E.A. and C.N. are supported by the National Institutes of Health grants R01HL15569 and R01HL144569.

## Author Contributions

C.N., C.A.C., and D.E.A. conceived and designed the experiments. J.N., P.W.W., T.N., C.N., C.A.C., and D.E.A. acquired, analyzed, and interpreted the data. J.N., P.W.W., and T.N. were responsible for the drafting of the manuscript. C.N., E.H.S., C.A.C., and D.E.A. critically revised the manuscript for important intellectual content. J.N., T.N., and C.A.C., performed statistical analysis. C.A.C. and D.E.A. obtained funding, provided administrative, technical, and material support. All authors read and approved the final manuscript.

## Abbreviations

CA: carbonic anhydrase
CPM: counts per million
DMR: differential methylated region
EWAS: epigenome-wide association study
GO: Gene Ontology
HEK293T: human embryonic kidney cell line
KEGG: Kyoto Encyclopedia of Genes and Genomes
KO: knockout
logFC: log fold-change
mtDNA: mitochondrial DNA
mtDNA-CN: mitochondrial DNA copy number
NC: normal control
nDNA: nuclear DNA
TFAM: mitochondrial transcription factor A
TMM: trimmed mean of M-values
ZNF: zinc finger proteins

## Supporting information

**S1 Fig. Top 20 Reactome pathways identified from differentially methylated sites (p-value < 1e-07).** N: number of genes identified in each term; ONT: ontology; BP: biological process; CC: cellular components; MF: molecular function.

**S2 Fig. Enriched KEGG pathways identified.** Red indicates hypermethylation/over-expressed genes; green indicates hypomethylation/under-expressed genes. A) DMSs – GABAergic synapse; B) DMRs – GABAergic synapse; C) DEGs – GABAergic synapse; D) DMSs – Protein Digestion and Absorption; E) DMRs – Protein Digestion and Absorption; F) DEGs – Protein Digestion and Absorption; G) DMSs – Neuroactive Ligand-Receptor Interaction; H) DMRs – Neuroactive Ligand-Receptor Interaction; I) DEGs – Neuroactive Ligand-Receptor Interaction.

**S3 Fig. Number of independent CpG sites associated with cancer or autoimmune disease (p < 0.05) in the EWAS catalog following 1000 randomized selections for the number of CpGs from the EWAS catalog identified in our primary analysis (785 CpGs).** Dashed line represents the actual number of significant CpGs from *TFAM* KO analysis that overlap reported CpGs from the EWAS catalog for each disease term. P-value represents enrichment for terms in our analysis.

**S1 Table. Summary statistics from identification of differentially methylated sites.**

**S2 Table. Summary statistics from identification of differentially methylated regions.** HMFDR: harmonic mean of the individual component FDRs

**S3 Table. Summary statistics from identification of differentially expressed genes.** LogFC: log fold change; logCPM: log counts per million; LR: likelihood ratio.

**S4 Table. Gene-CpG pairs discovered to be associated (negative or positive correlation).** Log2FC_Experiment.vs.Control: log2 fold change between the *TFAM* KO group and the NC group; Experiment.vs.Control.diff.pvalue: p-value of fold change difference between the *TFAM* KO group and the NC group.

**S5 Table. Overrepresented gene ontology terms identified from differentially methylated sites (p < 1e-07).** BP: biological process; MF: molecular feature; CC: cellular component; N: number of genes in GO term; DE: number of gene differentially methylated.

**S6 Table. Overrepresented gene ontology terms identified from differentially methylated regions (p < 1.17e-5).** BP: biological process; MF: molecular feature; CC: cellular component; N: number of genes in GO term; DE: number of gene differentially methylated.

**S7 Table. Overrepresented gene ontology terms identified from differentially expressed genes (p < 3.53e-6).** BP: biological process; MF: molecular feature; CC: cellular component; N: number of genes in GO term; DE: number of gene differentially methylated.

**S8 Table. Overrepresented KEGG pathways identified from differentially methylated sites (p < 1e-07).** N: number of genes in KEGG term; DE: number of gene differentially methylated.

**S9 Table. Overrepresented KEGG pathways identified from differentially methylated regions (p < 1.17e-5).** N: number of genes in KEGG term; DE: number of gene differentially methylated.

**S10 Table. Overrepresented KEGG pathways identified from differentially expressed genes (p < 3.53e-6).** N: number of genes in KEGG term; DE: number of gene differentially methylated.

**S11 Table. Overrepresented Reactome pathways identified from differentially methylated sites (p < 1e-07).** BP: biological process; MF: molecular feature; CC: cellular component; N: number of genes in Reactome term; DE: number of gene differentially methylated.

**S12 Table. Transcription factors identified from Fisher’s combined test of differentially methylated sites and differentially expressed genes.**

**S13 Table. Methylation sites and genes identified in the MitoCarta 3.0 database.** A) Methylation sites identified in the MitoCarta 3.0 database. B) Genes identified in the MitoCarta 3.0 database.

**S14 Table. Differentially expressed genes identified in the MitoCarta 3.0 database.**

**S15 Table. Differentially methylated sites overlapping with those reported in the EWAS Catalog.**

**S16 Table. Odds ratio (OR) and FDR for functional region enrichment.**

