## Supplementary figures and images for "Mitochondrial DNA copy number reduction via *in vitro TFAM* knockout remodels the nuclear epigenome and transcriptome"

### Supplemental Figure 1

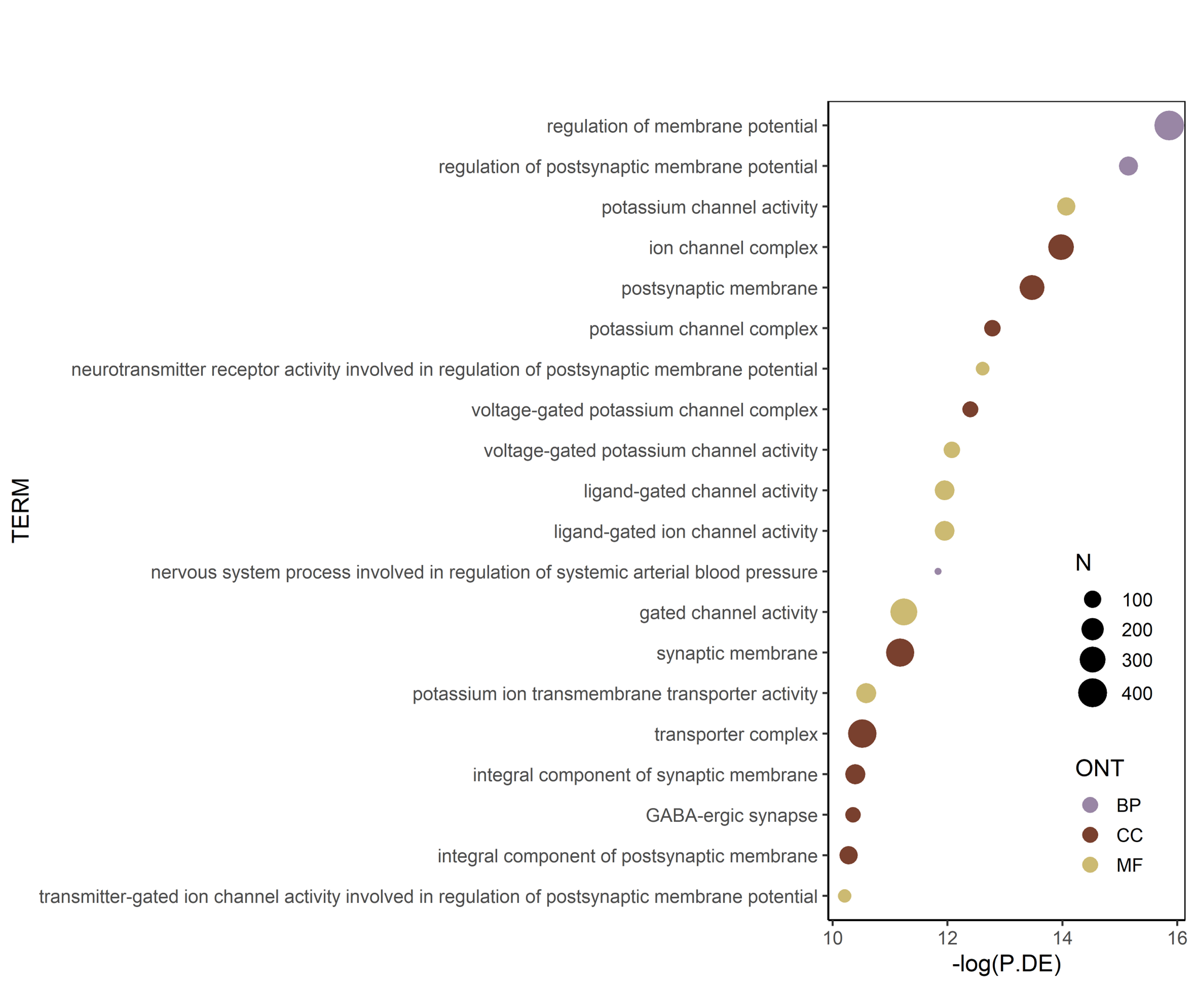

### Supplemental Figure 2

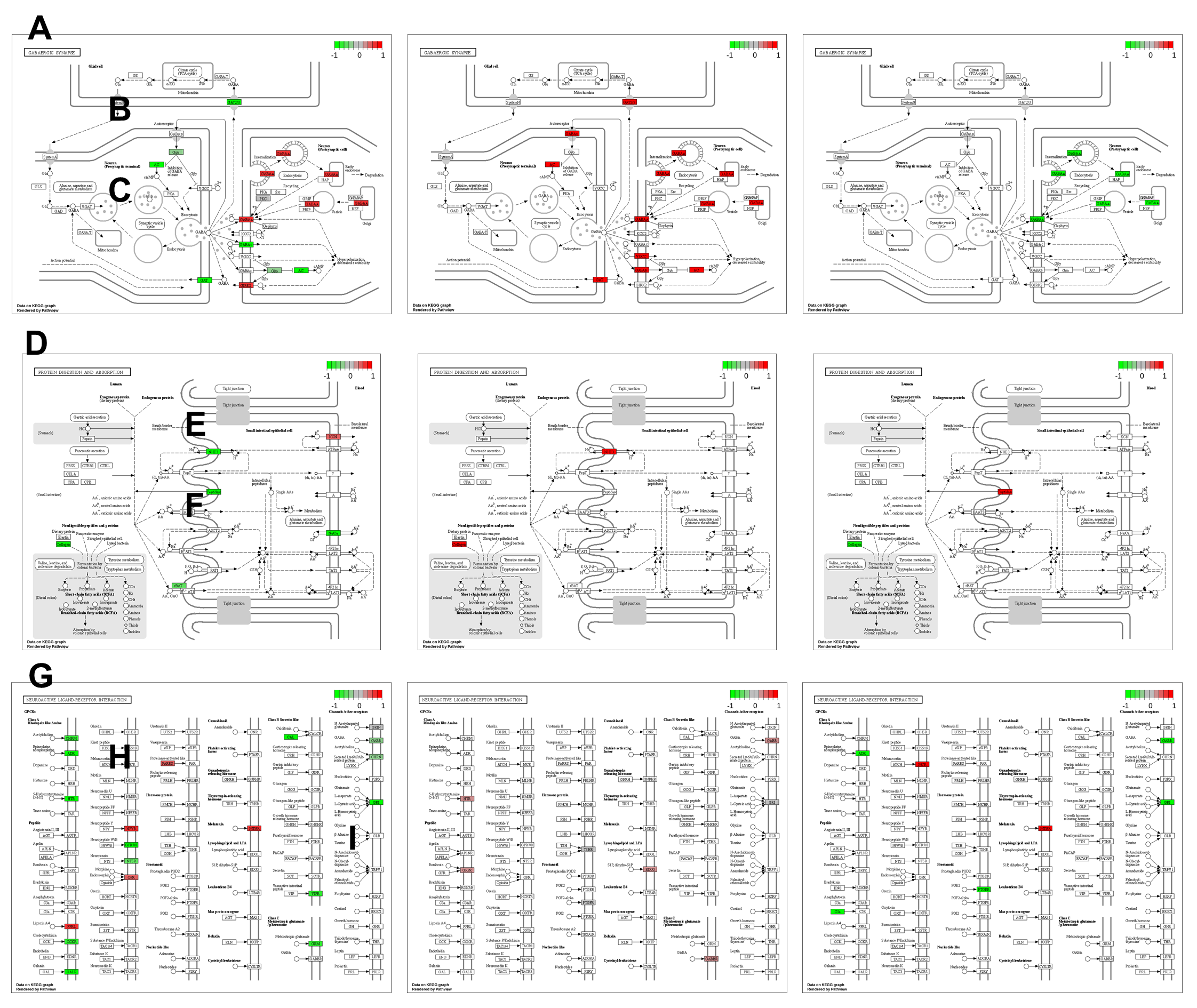

### Supplemental Figure 3

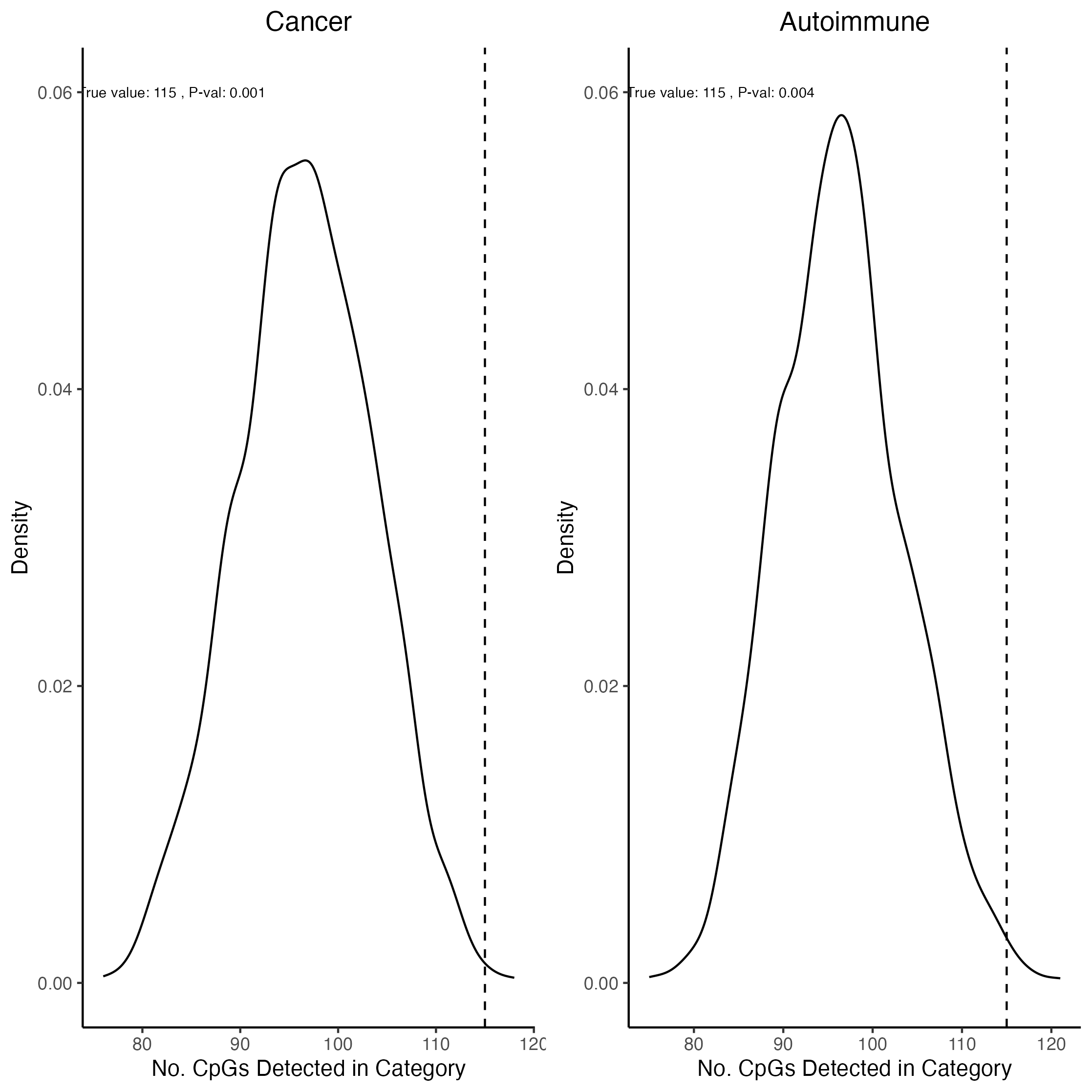
